# Toxicity assessment of 3-*O*-[6-deoxy-3-*O*-methyl-β-D-allopyranosyl-(1→4)-β-D-oleandropyranosyl]-17β–marsdenin isolated from *Gongronema latifolium* leaf on selected brain and kidney function indices in mice

**DOI:** 10.1101/2024.04.18.590095

**Authors:** Onyedika Gabriel Ani, Oluwatobi Ayodeji Medayedupin, Aminat Abike Azeez, Gideon Ampoma Gyebi, Isaac Duah Boateng, Joseph Oluwatope Adebayo

## Abstract

The safety of bioactive compounds, especially those isolated from medicinal plants, is a major concern for health authorities, pharmaceutical industries, and the public. Of recent, anti-tumor pregnane glycosides were isolated from *Gongronema latifolium* leaf, of which the toxicity of one, 3-*O*-[6-deoxy-3-*O*-methyl-β-D-allopyranosyl-(1→4)-β-D-oleandropyranosyl]-17β–marsdenin (3DMAOM), has not been evaluated. This study, therefore, evaluated the effects of 3DMAOM on selected brain and kidney function indices in mice. Female Swiss albino mice were randomly administered 5% dimethyl sulphoxide and different doses of 3DMAOM (0.5, 1, 2, and 4 mg/kg body weight) for fourteen (14) days, and their blood, brains, and kidneys were collected for biochemical analysis. There was no significant alteration in the activities of alkaline phosphatase (ALP), acetylcholinesterase, creatine kinase, Na^+^/K^+^-ATPase, Ca^2+^/Mg^2+^-ATPase, and Mg^2+^-ATPase in the brain of the treated groups compared to control. Also, no significant changes in the activities of ALP, gamma-glutamyltransferase, Na^+^/K^+^-ATPase, Ca^2+^/Mg^2+^-ATPase, and Mg^2+^-ATPase in the kidney of the treated groups compared to control. The plasma concentrations of Na^+^, K^+^, Cl^-^, PO_4_^3-^, creatinine, and urea of mice were not significantly altered at all doses of the 3DMAOM compared to controls. However, the plasma concentration of Ca^2+^ was significantly reduced (p<0.05) at all doses of the 3DMAOM, and the plasma concentration of uric acid was significantly reduced (p<0.05) at 2 mg/kg body weight of the 3DMAOM compared to controls. These findings suggest that 3DMAOM isolated from *Gongronema latifolium* leaf may not adversely affect brain function but may affect calcium ion homeostasis in subjects.

## 1. Introduction

Natural products (secondary metabolites) have been well-documented as therapeutic agents. They have been the most successful sources of new and approved drugs for treating human diseases, as well as a source of leads for potential drug discovery [1]. Plants afford structurally distinct secondary metabolites known as phytochemicals with diverse pharmacological properties [2–3], thus making them primary sources of modern pharmaceuticals used for various ailments [4]. These phytochemicals are mainly non-nutrient compounds in various parts of plants, such as fruits, roots, stem bark, and leaves, some of which can potentially be used to treat diverse diseases. Among these phytochemicals, the most abundant include alkaloids, flavonoids, saponins, sterols, phenolic compounds, and glycosides [5]. Of recent, some anti-cancer pregnane glycosides were isolated from *Gongronema latifolium* [6].

*G. latifolium* Benth (**Figure 1**), formerly *Marsdenia latifolia*, is a tropical rainforest herbaceous climber belonging to the family of *Asclepiadacea* that produces a yellow flower and a characteristic milky exudate when the stem is incised [6]. *G. latifolium* is commonly known as amaranth globe, in English, ‘utazi’ and ‘arokeke’ or ‘madumaro’ in South-Eastern and South-Western Nigeria, respectively, where it is used as a spice and vegetable [7]. In Southeast Nigeria, the leaves of *G. latifolium* are used to prepare food for puerperal mothers. It is claimed to arouse appetite, diminish post-partum contraction, and stimulate normalcy of menstrual cycle hormones [8] and spice locally brewed beer [7]. *G. latifolium* leaf has been reported to have antihyperglycemic, anti-lipidemic, anti-hypercholesterolemic, and antioxidant effects [9–11]. It also has antimicrobial, antitussive, and antidiarrheal properties [12]. The ulcer-inhibitory and hepatoprotective effects of the plant have also been reported [13–14].

**Figure 1:**
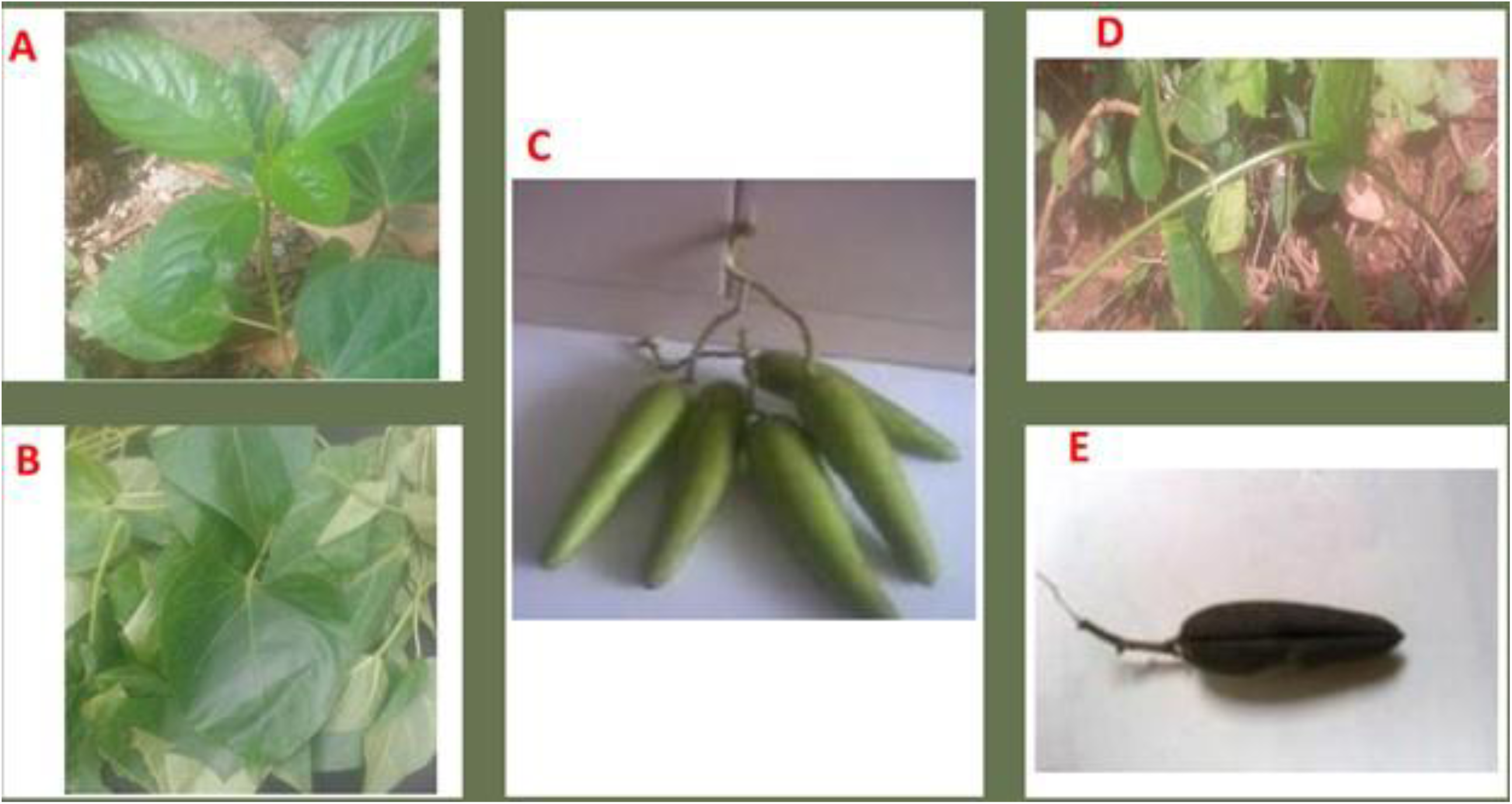
Tender Leaves (A and B) Bunch of fleshy fruits (C) Soft and pliable stem (D) and dry fruit (E) of *G. latifolium*

Phytochemical screenings on the fruits and leaves of *G. latifolium* identified the presence of alkaloids, tannins, saponins, flavonoids, phytic acid, hydrocyanic acid, phenols, lignan, terpenes, sterol, allicin, hydroxycinnamic acids, and carotenoid, with some in appreciable concentrations in the leaf of the plant [15–16]. Minerals (Cr, Cu, Se, Zn, and Fe) and vitamins (A, C, E, riboflavin, thiamine, and niacin) are also present in the plant’s root, bark, and twig extracts [17]. The abundance of phytochemicals and micronutrients in *Gongronema latifolium* confers nutritional, industrial, therapeutic, and economic potential. Chemical investigation of the leaf extract led to the isolation of a novel pregnane glycoside: iloneoside (3-*O*-[6-deoxy-3-*O*-methyl-β-D-allopyranosyl-(1→14)-β-D-oleandropyranosyl]-11,12-di-*O*-tigloyl-17β-marsdenin), together with two already known pregnane glycosides, 3-*O*-[6-deoxy-3-*O*-methyl-β-D-allopyranosyl-(1→4)-β-D-oleandropyranosyl]-17β-marsdenin (3DMAOM) (**Figure 2**) and 3-*O*-[6-deoxy-3-*O*-methyl-β-D-allopyranosyl-(1→4)-β-D-canaropyranosyl]-17β-marsdenin. These pregnane glycosides elicited *in vitro* anti-proliferative effects, with iloneoside being the most active [6].

**Figure 2:**
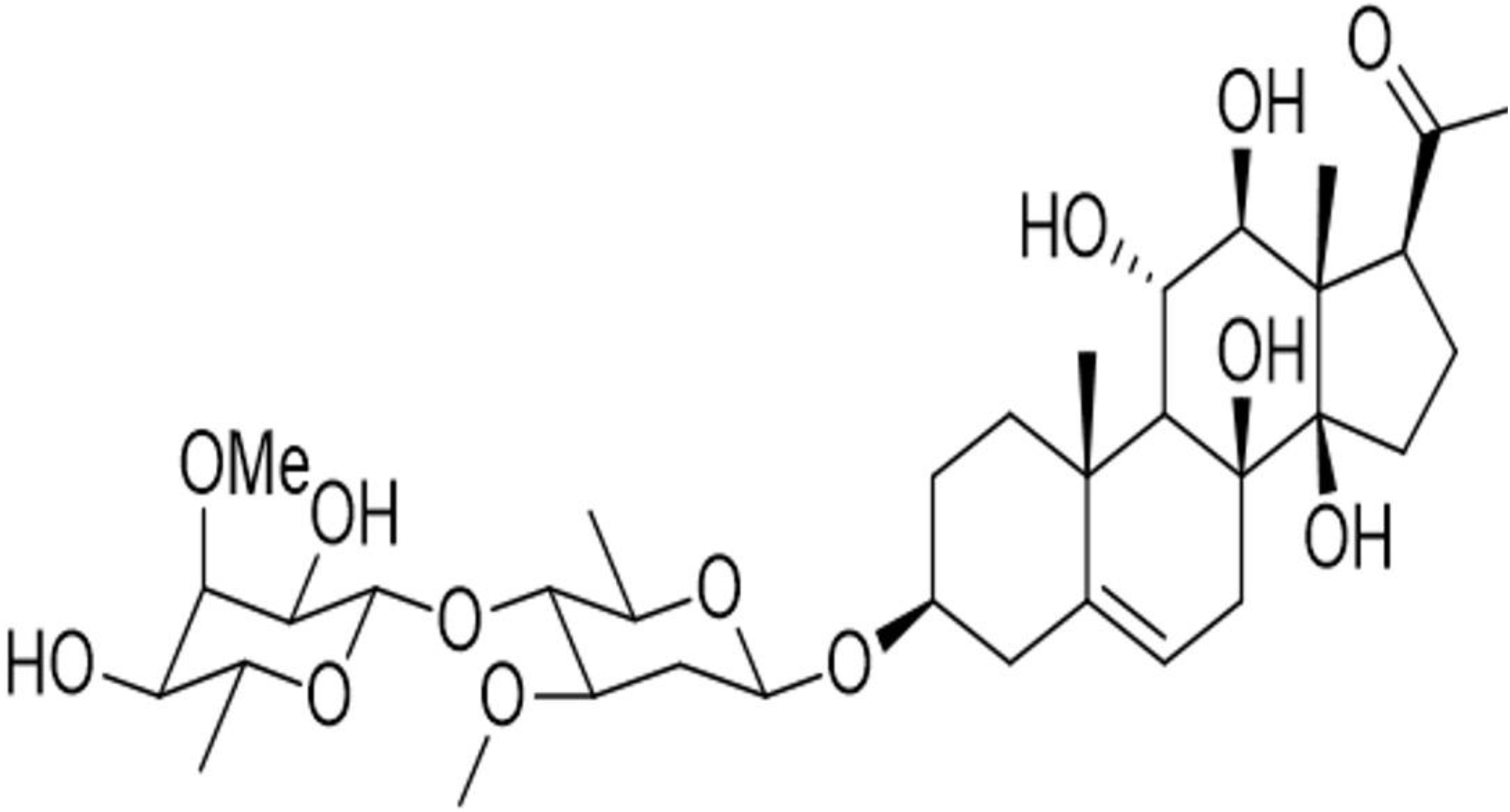
3-*O*-[6-deoxy-3-*O*-methyl-β-D-allopyranosyl-(1→4)-β-D-canaropyranosyl]-17β-marsdenin.(adapted from [6] with permission)

Pregnanes and their glycosides have attracted considerable attention recently because of their cytotoxic [6–18] and anti-proliferative activities [19–21]. Also, they have been reported to exhibit antifungal [22], anti-herpes [20], and appetite suppressor activities [23–24]. However, the risk of toxicity associated with using natural products is one of the main reasons for the hesitance among healthcare practitioners towards promoting their integration into healthcare systems. Recently, high doses of 3-*O*-[6-deoxy-3-*O*-methyl-β-D-allopyranosyl-(1→4)-β-D-canaropyranosyl]-17β-marsdenin were associated with cardiac muscle adverse effects [25]. Therefore, there is a need to evaluate the toxicity of the other isolated pregnane glycosides from *G. latifolium* leaf.

This study, therefore, evaluated the effects of 3DMAOM on selected brain and kidney function indices in mice. The critical role of the brain in central physiology and the kidney’s homostatic function warrants the investigation of any potential side effects such agents may pose. Pharmacological side effects are common and have resulted in the withdrawal of many potent agents, especially in cancer chemotherapy. Therefore, it is pertinent despite the reported anticancer potential of the pregnane glycosides, their safety should also be evaluated.

## 2. Materials and methods

### 2.1. Chemical and reagents

Absolute methanol and ethanol were purchased from the British Drug House, Poole, England. Diethyl ether was obtained from May and Baker Ltd., Dagenham, England, while Dimethyl sulphoxide was obtained from Sigma-Aldrich Chemical Company, St. Louis, Mo, USA. Assay kits were purchased from Randox Laboratories Ltd., Co-Antrim, UK. All other reagents and equipment were of analytical grade without further purification.

### 2.2. Pregnane glycoside

The pregnane glycoside, 3DMAOM (**Figure 2**), was isolated in a previous study [6].

### 2.3. Experimental animals

Twenty female albino mice (*Mus musculus*) of an average weight of 18 ± 2 g were obtained from the Small Animal Holding Unit of the Department of Biochemistry, Faculty of Life Sciences, University of Ilorin, Ilorin, Nigeria. The mice were acclimatized for fourteen days under good laboratory conditions (12 h light/dark cycle; room temperature: 28 - 30°C). They were allowed access to standard rodent chow (Vital feed^®^, Grand Cereals Ltd, Jos, Nigeria) and water *ad libitum*. Handling, management, and use of animals were according to guidelines approved by the University Ethical Review Committee of the University of Ilorin, Ilorin, Nigeria. At the end of the fourteen-day acclimatization period, the animals were randomly assigned to five groups of ten mice each. The 3DMAOM dissolved in 5% DMSO was orally administered to the mice with the aid of an esophageal cannula as follows:

**Group A:** 5% DMSO solution

**Group B:** 0.5 mg/kg body weight of the 3DMAOM

**Group C:** 1 mg/kg body weight of the 3DMAOM

**Group D:** 2 mg/kg body weight of the 3DMAOM

**Group E:** 4 mg/kg body weight of the 3DMAOM

### 2.4. Determination of Organ-Body Weight Ratio

The relative weight of the organ to the body weight of the corresponding mouse, expressed as a percentage, was calculated using the formula:

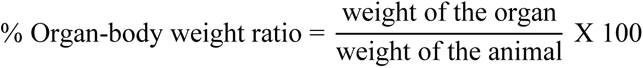

### 2.5. Biochemical analysis

#### 2.5.1. Sample collection and preparation

Twenty-four hours after the last set of doses was administered, the mice were sacrificed under mild diethyl ether anesthesia. The neck region was cleared of fur, and venous blood was collected into lithium heparin sample bottles. The heparinized blood samples were centrifuged at 3,000 rpm for 5 min, and the plasma was carefully pipetted into properly labeled tubes. These were stored frozen until needed for analysis. Organs of each mouse (brain and kidney) were excised, cleansed of superficial connective tissues, weighed, and homogenized in ice-cold 0.25 M sucrose solution (1:5 w/v). These were centrifuged at 1,000 rpm for 5 min in a centrifuge, and the supernatants were pipetted into new sample tubes. These were stored at -20°C overnight to ensure the maximum release of enzymes. The supernatant was used for the analysis below.

#### 2.5.2. Total protein concentration

The total protein concentrations in the animals’ brains, kidneys, and serum were assayed using Biuret reagent, as described by Gornall *et al.* [26]. Biuret reagent (4.0 ml) was added to 1.0 ml of the sample. This was mixed thoroughly by shaking and left undisturbed for 30 minutes at room temperature for color development. The blank was constituted by replacing the sample with 1.0 ml of distilled water. The absorbance was read against blank at 540 nm using a UV-VIS spectrophotometer. The protein concentration in the sample was calculated by comparing them with those on the calibration curve for egg albumin. The concentration of the protein in the sample was extrapolated from the calibration curve of the egg albumin using the expression:

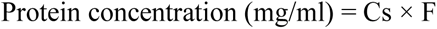

Where:Cs = corresponding protein concentration from the calibration, F = dilution factor

#### 2.5.3. Alkaline phosphatase activity

The method described by Wright *et al.* [27] was employed to assay the alkaline phosphatase activities in the brain and kidney. In series, 2.2 ml of carbonate buffer (0.1 M) and 0.1 ml of MgSO_4_.7H_2_O (0.1 M) were added to the test tubes. Then 0.2 ml of the enzyme source was added and incubated at 37 °C. for 10 minutes. 0.5 ml of 10 mM p-nitrophenyl phosphate (substrate) was added, and the assay mixture was incubated again for 30 minutes at 37°C. The reaction was terminated immediately by adding 2.0 ml of 1N sodium hydroxide. The blank was constituted by replacing the enzyme source with 0.2 ml of distilled water. The absorbance was read spectrophotometrically at 400 nm, and the results were expressed as U/mg protein.

#### 2.5.4. Gamma-Glutamyltransferase activity

Gamma-glutamyltransferase (GGT) assay was done using the procedure described by Szasz *et al.* [28]. 1ml of the reconstituted reagent (comprising of 3ml of buffer/Glycineglycyl and one vial of L-ɤ-glutamyl-3-carboxy-4-nitroanilide) was added to 0.1ml of sample. The blank was constituted by replacing the sample with distilled water. The mixture was mixed properly, and initial absorbance was read at 405nm immediately and at 1, 2, and 3 minutes. The activity of GGT was calculated as follows and expressed as U/mg protein.

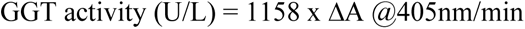

#### 2.5.5. Creatine kinase-MB activity

The method described by Di Witt and Trendelendurg [29] was used for assaying the activity of creatine kinase-MB. 1000 µl of working reagent was added to 40 µl of the sample, it was incubated at 37°C for 100 seconds. Change in absorbance was then read at 340nm per minute for 5 minutes and expressed as U/mg protein.

#### 2.5.6. Acetylcholinesterase (AChE) activity

Acetylcholinesterase (AChE) activity was assayed using the method described by Magnotti *et al.* [30]. To separate test tubes labeled blank and standard, 100 μl of distilled water and 100 μl of calibrator (20 U/L) were added. Aliquot (50 μl) of water and 100 μl of calibrator was incubated at room temperature in a final reaction mixture of 1 ml, containing 100 nm phosphate buffer (pH 7.5), 1 nM 5,5^’^-dithiobisnitrobenzoic acid and 0.8 nm acetylthiocholine iodide. After 2 and 10 min, initial and final sample absorbances, at 412 nm, were read and expressed as U/mg protein.

#### 2.5.7. ATPase activity

The determination of Ca^2+^ and Mg^2+^-ATPase activity was carried out using the method of Bewaji *et al*. [31]. Appropriately diluted 10µl of the sample was added to a test tube already containing 500µl of Ca^2+^ ATPase buffer (120mM KCl/30mM Tris, pH 7.4), 20µl of 20mM CaCl_2_, 10µl of MgCl_2_. Distilled water was further added to make the total reaction volume 800µl with 100µl ATP (substrate), which was used to initiate the reaction after incubating at 37°C for 3 minutes after the reaction was initiated by adding ATP, it was incubated at 37°C for 30 minutes, after which the reaction was stopped using 200µl 5% Sodium Dodecyl Sulfate (SDS). Finally, 2ml of reagent C was added and left at room temperature for color development, after which the absorbance was read at 820nm. Note that the reaction was carried out in triplicate for each sample and that the volume of ATP was varied from 100µl to 10µl to get the reaction kinetics. The ATPases were expressed as mmol/min/mg protein.

The determination of Mg^2+^-ATPase activity was carried out using the method described by Fleschner and Kraus-Friedmann [32]. For the test, 400 µl of 240 mM KCl / 60 mM Tris (pH 7.4) was pipetted into the test tube. Thereafter, 20 µl of MgCl_2_.6H_2_O (80 mM), 20µl of EGTA (20 mM), 220 µl of distilled water, 20 µl of appropriately diluted enzyme source, and 20 µl of Vanadate (0.04 M) was added. This was mixed and incubated at 37°C for 5 minutes. Then 100µl of ATP (8 mM) was added, this was also mixed and incubated at 37°C for 30 minutes. Thereafter, 200 µl of SDS (5%) and 2 ml of reagent C were added. The mixture was left undisturbed at room temperature for 30 minutes for color development. The blank was similarly prepared, but 20 µl of distilled water was used instead of the 20 µl enzyme source. The absorbance of the test was read against the blank at 820 nm. The absorbance obtained was then extrapolated from the calibration curve for phosphate to obtain inorganic phosphate concentration.

Assay for Na^+^, K^+^-ATPase activity was estimated according to Bewaji *et al.* [31]. 400µl of the buffer (200mM NaCl/40mM, KCl/60mM, Tris, pH 7.4), 20µl tissue sample mixed in that order and incubated at 370C minutes. Then, 200µl of (5%) SDS 2000 of reagent C was added, and the mixture was left at room temperature for half an hour for color development. In the blank, 20µl of distilled water replaced 20µl of tissue sample. The absorbance of the test was read at 820nm. The absorbance obtained was then extrapolated from the calibration curve for phosphate to obtain the inorganic phosphate concentration.

#### 2.5.8. Plasma concentrations of mineral ions

Flame emission photometry was used to determine plasma sodium and potassium ion concentration. Aliquot (0.1 mL) of plasma was diluted with 19.9 mL of distilled water and shaken. The samples were then aspirated into a flame photometer, and the results were read (in mmol/L on a digital read-out device) after setting the instrument to zero with glass distilled water and calibrated with respective standards (140 mEq/L for Na^+^ and 5 mEq/L for K^+^).

Calcium ion concentration in plasma was determined by the method described by Biggs and Moorehead [33]. 1000 μL of the working reagent (360 mmol/L diethylamine, 0.15 mmol/L O-cresolphalein complex, 17.2 mmol/8-hydroquinoline) was added to separate test tubes containing aliquot (10 μL) of sample, standard calcium solution (10 mg/dL) and distilled water which serves as test, standard and blank respectively. These were mixed and incubated at room temperature for 5 min. The absorbances of the test (sample) and standard were read against the reagent blank at a wavelength of 578 nm.

The concentration of chloride ions was determined by the method described by Yoshinaga and Ohta [34]. Aliquot (1000 μL) of the working reagent (2 mmol/L mercuric (II) thiocyanate, 29 mmol/L nitric acid, 20 mmol/L ferric nitrates) was added to separate test tubes containing Aliquot (10 μl) of the sample, standard solution and distilled water which served as test, standard and blank respectively. These were mixed and incubated at 37°C for 1 min. The absorbance of the sample and standard were read against the reagent blank at 492nm.

Phosphate ion concentration was determined using the method described by Tietz [35]. Aliquot (1000 μL) of the working reagent (210 mmol/L sulphuric acid, 650 mmol/L ammonium molybdite) was pipetted into test tubes for blank, standard, and sample, respectively. Then 20 μL of distilled water, standard solution (5 mg/dL), and sample were pipetted into the respective test tubes, mixed, and incubated at 37°C for 1 min. The absorbances of standard and tests were read against reagent blank at 340 nm.

#### 2.5.9. Plasma creatinine concentration

Plasma creatinine concentrations were determined using the method described by Bartels and Bohmer [36]. 1.0 ml of the working reagent was added to 0.1 ml of the sample. The standard was constituted by replacing the sample with 0.1 ml of the standard solution. The solution was mixed, and after 30 seconds, the absorbance A1 of the standard and sample was read. Exactly 2 minutes later, absorbance A2 of standard and sample was read. Absorbance was read at 492 nm against air.

#### 2.5.10. Plasma uric acid and urea concentrations

Plasma uric acid was determined according to the method described by Tietz [35]. 1 ml of the reagent was added to test tubes containing 0.02 ml of serum. The blank and standard were constituted by replacing the serum with 0.02 ml of distilled water and standard working reagent, respectively. The mixture was incubated for 5 minutes at 37°C, and the absorbance of the sample and standard were read spectrophotometrically against a reagent blank within 30 minutes at a wavelength of 520 nm.

The urea concentration in the plasma was determined using the Urease-Berthlot method, as described by Weatherburn [37]. 0.1 ml of reagent 1 was added to test tubes containing 0.01 ml of serum (appropriately diluted) blank and standard. The blank and standard were constituted by replacing the serum with 0.01 ml of distilled water and standard working reagent. The reaction constituent was thoroughly mixed and incubated at 37^0^C for 10 minutes. 2.5ml of reagents 2 and 3 were added sequentially to the test tubes (sample, blank, and standard). The reaction constituent was thoroughly mixed and incubated again at 37^0^C for 15 minutes. The absorbance of the sample and standard were read against the blank spectrophotometrically at 546 nm.

### 2.6. Statistical analysis of data

All experiments were done in four replicates, and the results were expressed as mean ± standard error of the mean. The one-way analysis of variance (ANOVA) was used to analyze the data for statistical significance, followed by Turkey’s post hoc multiple comparisons, using IBM SPSS Statistics for Windows, Version 23.0 (IBM Corp., Armonk, N. Y., USA). Difference at P<0.05 was considered significant. Graphs were created using GraphPad Prism 7 software for Windows (GraphPad Software, California, USA). Origin 2019b software (OriginLab^®^, MA, USA) was used for hierarchical cluster analysis and principal component analysis,

## 3. Results

### 3.1. The organ-body weight ratios were unaltered following the administration of 3DMAOM

The organ’s relative weight to the corresponding mouse’s body weight was assessed, as shown in **Table 1**. The results revealed that the change in mice’s kidney- and brain-body weight ratios were not statistically significantly different at all doses of the 3DMAOM administered compared to controls.

**Table 1:** Effects of 3-O-[6-deoxy-3-O-methyl-β-D-allopyranosyl-(1→4)-β-D-oleandropyranosyl] -17β–marsdenin on selected organ-body weight ratios of mice.

| Treatment | Organ-Body weight ratio (%) |  |
| --- | --- | --- |
|  | Brain | Kidney |
| 5% DMSO | 1.77 ± 0.20<sup>a</sup> | 1.20 ± 0.07<sup>a</sup> |
| 0.5 mg/kg bwt 3DMAOM | 2.18 ± 0.32<sup>a</sup> | 1.32 ± 0.08<sup>a</sup> |
| 1 mg/kg bwt 3DMAOM | 2.13 ± 0.06<sup>a</sup> | 1.31 ± 0.14<sup>a</sup> |
| 2 mg/kg bwt 3DMAOM | 2.08 ± 0.34<sup>a</sup> | 1.38 ± 0.15<sup>a</sup> |
| 4 mg/kg bwt 3DMAOM | 1.78 ± 0.23<sup>a</sup> | 1.23 ± 0.01<sup>a</sup> |

Values are Means ± SEM of 4 replicates. Values in the same column with different superscripts are significantly different (p<0.05).
DMSO = Dimethyl sulphoxide; bwt = body weight; 3DMAOM= 3-*O*-[6-deoxy-3-*O*-methyl- β-D-allopyranosyl-(1 → 4)- β-D- oleandropyranosyl]-17 $\beta$ - marsdenin.

### 3.2 Administration of 3DMAOM did not alter the activities of membrane-bound and cellular enzymes in both the brain and kidney of mice

There was no significant alteration in alkaline phosphatase (ALP) activities in the brain and kidney of mice administered various doses of the 3DMAOM compared to controls (**Figure 3A**). The administration of the 3DMAOM at various doses had no significant effect on Ca^2+^/Mg^2+^-ATPase, Mg^2+^-ATPase, Na^+^/K^+^-ATPase activities in the brain and kidneys of mice compared to the controls (**Figures 3B-D and 3B**). In the kidney, GGT activities did not significantly differ in any of the doses compared to the control **(Figure 4**). Also, creatine kinase (CK) and AChE activities in the brain of mice were not significantly changed at all doses of 3DMAOM administered compared to controls (**Figures 4A and 4C**, respectively).

**Figure 3:**
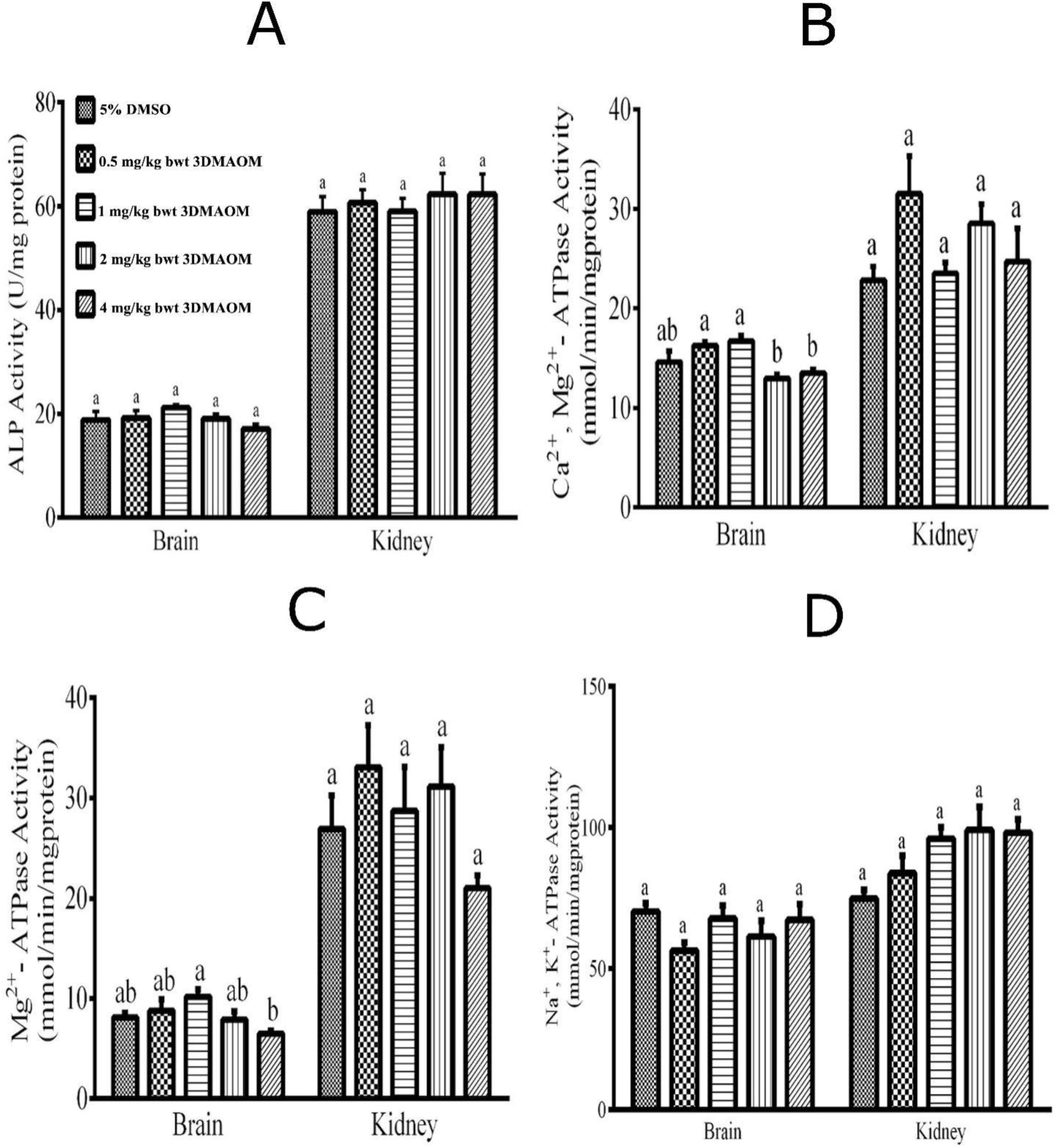
Effects of 3-*O*-[6-deoxy-3-*O*-methyl-β-D-allopyranosyl-(1→4)-β-D-oleandropyranosyl] -17β–marsdenin on activities of selected membrane-bound enzymes in brain and kidney of mice.(A) ALP activity (B) Ca^2+^, Mg^2+^-ATPase (C) Mg^2+^-ATPase (D) Na^+^, K^+^-ATPase Values are Means ± SEM of 4 replicates. Bars for each organ with a different alphabet indicated significant difference (p<0.05). **Note.** DMSO = Dimethyl sulphoxide; bwt = body weight; 3DMAOM = 3-O-[6-deoxy-3-O-methyl-β-D-allopyranosyl-(1→4)-β-D-oleandropyranosyl]-17β–marsdenin; ALP = Alkaline phosphatase.

**Figure 4:**
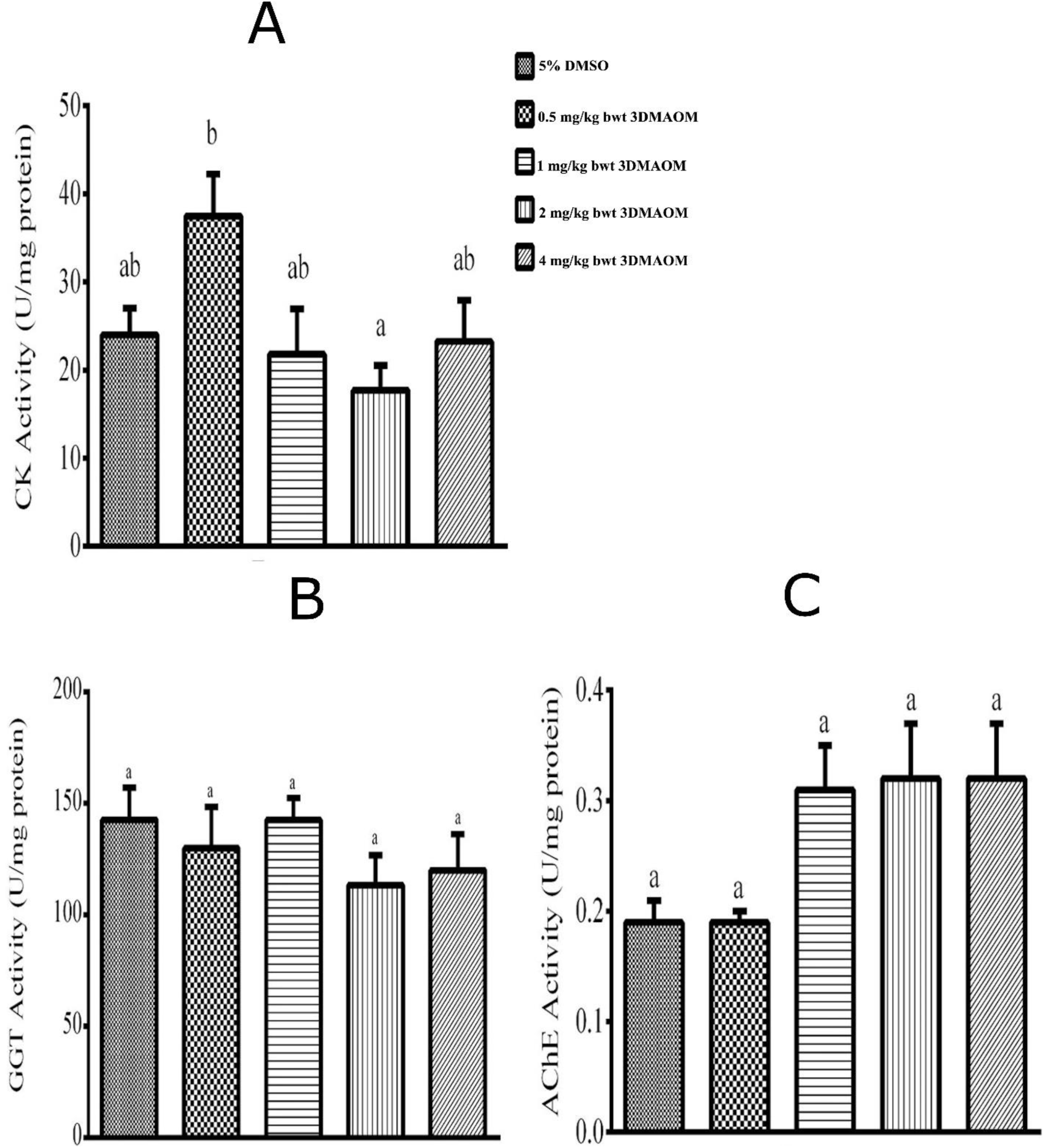
Effects of 3-*O*-[6-deoxy-3-*O*-methyl-β-D-allopyranosyl-(1→4)-β-D-oleandropyranosyl] -17β–marsdenin on specific activities of selected cellular enzymes of brain and kidney of mice. **(A)** CK **(B)** GGT **(C)** AChE Values are Means ± SEM of 4 replicates. Bars for each organ with a different alphabet indicated significant difference (p<0.05). **Note.** DMSO = Dimethyl sulphoxide; bwt = body weight; 3DMAOM = 3-O-[6-deoxy-3-O-methyl-β-D-allopyranosyl-(1→4)-β-D-oleandropyranosyl]-17β–marsdenin; CK = creatine kinase; GGT = gamma-glutamyltransferase; AChE = Acetylcholinesterase.

### 3.3 3DMAOM administration did not affect the plasma concentrations of electrolytes and biomolecules in the mice, except plasma Ca^2+^ and uric acid concentrations

The administration of 3DMAOM at various doses had no significant effect on plasma concentrations of Na^+^, K^+^, Cl^-^, and PO_4_^3-^ compared to controls (**Table 2**). However, there was a significant reduction (p<0.05) in the plasma concentration of Ca^2+^ at all doses of 3DMAOM compared to the control (**Table 2**). Administration of 3DMAOM at all doses had no significant effect on the plasma concentrations of creatinine and urea compared to the controls (**Table 3**). However, the administration of 3DMAOM at 2 mg/kg body weight caused a significant reduction (p<0.05) in plasma concentration of uric acid, with no significant alteration at other doses compared to control (**Table 3**).

**Table 2:** Effects of 3-O-[6-deoxy-3-O-methyl-β-D-allopyranosyl-(1→4)-β-D-oleandropyranosyl]-17β–marsdenin on plasma concentrations of selected electrolytes of mice.

| Treatment | Na<sup>+</sup> (mEq/dL) | Ca<sup>2+</sup> (mEq/dL) | K<sup>+</sup> (mEq/dL) | Cl<sup>-</sup> (mEq/dL) | PO<sub>4</sub><sup>3-</sup> (mEq/dL) |
| --- | --- | --- | --- | --- | --- |
| 5% DMSO | 67.20± 1.27<sup>a</sup> | 20.18± 0.69<sup>a</sup> | 12.11± 1.74<sup>a</sup> | 39.53± 0.42<sup>a</sup> | 4.85± 0.93<sup>a</sup> |
| 0.5mg/kg bwt 3DMAOM | 70.79± 3.32<sup>a</sup> | **14.71± 1.24<sup>b</sup>** | 8.98± 1.10<sup>a</sup> | 38.21± 0.34<sup>a</sup> | 4.03± 0.25<sup>a</sup> |
| 1mg/kg bwt 3DMAOM | 65.97± 1.44<sup>a</sup> | **10.81± 0.90<sup>c</sup>** | 8.64± 0.99<sup>a</sup> | 38.88± 0.78<sup>a</sup> | 4.78± 0.24<sup>a</sup> |
| 2mg/kg bwt 3DMAOM | 63.46± 1.33<sup>a</sup> | **10.30± 0.11<sup>c</sup>** | 9.06± 0.86<sup>a</sup> | 38.63± 0.49<sup>a</sup> | 5.46± 0.45<sup>a</sup> |
| 4mg/kg bwt 3DMAOM | 64.50± 1.21<sup>a</sup> | **9.02± 0.56<sup>c</sup>** | 7.52± 0.26<sup>a</sup> | 38.56± 0.49<sup>a</sup> | 6.49± 1.21<sup>a</sup> |

Values are Means ± SEM of 4 replicates. Values in the same column with different superscripts are significantly different (p<0.05).
**Note.**
DMSO = Dimethyl sulphoxide; bwt = body weight; 3DMAOM = 3-O-[6-deoxy-3-O-methyl-β-D-allopyranosyl-(1 → 4)- β-D-oleandropyranosyl]-17 β-marsdenin.

**Table 3:**
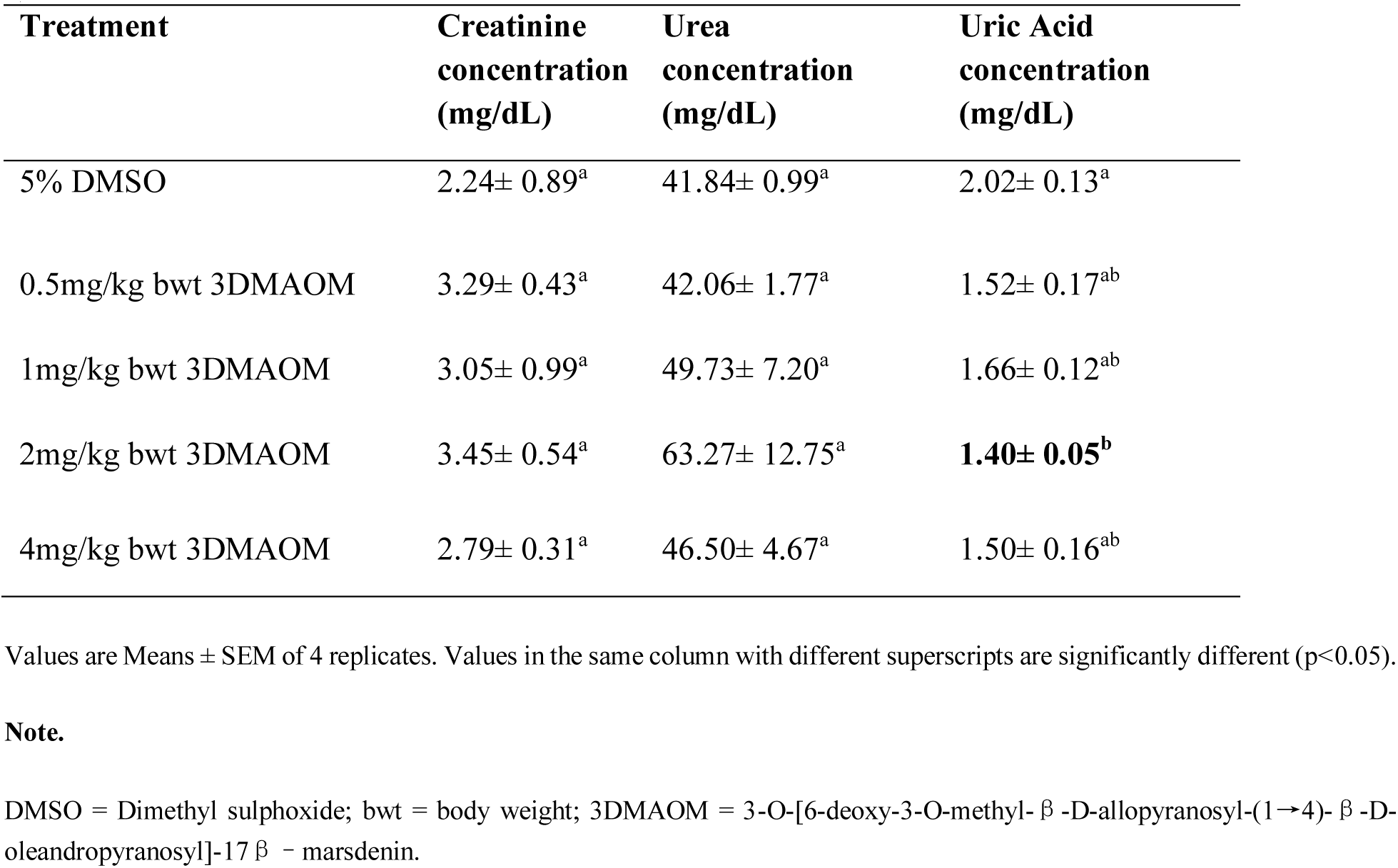
Selected kidney function indices of mice administered 3-*O*-[6-deoxy-3-*O*-methyl-β-D-allopyranosyl-(1→ 4)-β-D-oleandropyranosyl] -17β–marsdenin.

| **Treatment** | **Creatinine**<br>**concentration**<br>**(mg/dL)** | **Urea**<br>**concentration**<br>**(mg/dL)** | **Uric Acid**<br>**concentration**<br>**(mg/dL)** |
| --- | --- | --- | --- |
| 5% DMSO | 2.24± 0.89<sup>a</sup> | 41.84± 0.99<sup>a</sup> | 2.02± 0.13<sup>a</sup> |
| 0.5mg/kg bwt 3DMAOM | 3.29± 0.43<sup>a</sup> | 42.06± 1.77<sup>a</sup> | 1.52± 0.17<sup>ab</sup> |
| 1mg/kg bwt 3DMAOM | 3.05± 0.99<sup>a</sup> | 49.73± 7.20<sup>a</sup> | 1.66± 0.12<sup>ab</sup> |
| 2mg/kg bwt 3DMAOM | 3.45± 0.54<sup>a</sup> | 63.27± 12.75<sup>a</sup> | **1.40± 0.05<sup>b</sup>** |
| 4mg/kg bwt 3DMAOM | 2.79± 0.31<sup>a</sup> | 46.50± 4.67<sup>a</sup> | 1.50± 0.16<sup>ab</sup> |

Values are Means ± SEM of 4 replicates. Values in the same column with different superscripts are significantly different ( $p < 0.05$ ).
**Note.**
DMSO = Dimethyl sulphoxide; bwt = body weight; 3DMAOM = 3-*O*-[6-deoxy-3-*O*-methyl- β-D-allopyranosyl-(1 → 4)- β-D-oleandropyranosyl]-17 β-marsdenin.

## 4. Discussion

This study investigated the toxicity effects of 3DMAOM isolated for *G. latifolium* on crucial functions of the brain and kidney in mice. Aside from the interference of Ca^2+^ homeostasis and the production of uric acid (**Table 2**), the 3DMAOM did not produce any significant changes in either the brain or the kidney of mice, suggesting their relative safety. Fourteen days of administration of the 3DMAOM to mice did not significantly alter the weights of the brain and kidney relative to the mouse’s body weight. This suggests that the 3DMAOM may not cause constriction (resulting from necrosis) or inflammation of these organs. Weight changes could explain the cellular energy metabolism in animals including mice. Specifically, the organ-body weight ratios elucidate closely any interference in the organ’s energy pathways [38]. In toxicological studies, organ-body weight ratios are evaluated to determine the change in organ weight relative to the whole weight of the animal to identify potentially harmful effects of chemicals [39]. This organ-body weight ratio is often useful for ascertaining treatment-related organ weight changes, as significant differences between treated and control animals may occur without morphological changes. Several studies have shown that plant extracts can significantly impact the organ-body weight ratio in animal subjects [38–39]. Such changes could be due to the dosage or the duration of the administration. Factors other than the direct toxic effects of chemicals also cause changes in body weight and associated changes in organ weight by secondary means. In rodents and other species, the absolute weights of many, but not all, organs are affected by growth during the normal life span [42]. Logically, a chemical that alters body weight would not change the absolute weight of those organs that do not significantly change in weight during normal growth (e.g., brain or kidney) unless there was another specific toxic effect on the organ [39]. A major goal of a toxicity study is to identify target organs susceptible to toxic doses of the test compound.

Every major body system can be adversely affected by toxic substances, but the nervous system (the brain, the spinal cord, and a vast array of nerves and sensory organs that control major body functions) is particularly vulnerable [43]. The nervous system is influenced by the functioning of other organ systems (e.g., hepatic, cardiovascular, and endocrine systems). Thus, toxicant-induced alterations in any organ system can be reflected in changes in neurobehavioural output. This fact alone suggests that nervous system function should be among the first to be thoroughly assessed in cases of exposure to known or potentially hazardous agents. Many neurotoxic substances can cause death when absorbed, inhaled, or ingested in sufficiently large quantities. Neurotoxic substances play a significant causal role in the development of some neurological and psychiatric disorders [43]. A substance is considered to have neurotoxic potential if it adversely affects any of the structural or functional components of the nervous system. A substance might interfere with protein synthesis in certain nerve cells at the molecular level, reducing neurotransmitter production and brain dysfunction. At the cellular level, a substance might alter the flow of ions (charged molecules, e.g., sodium and potassium ions) across the cell membrane, thereby perturbing the transmission of impulses between nerve cells. Substances that adversely affect sensory or motor function disrupt learning and memory processes or cause detrimental behavioral effects are neurotoxic, even if the underlying molecular and cellular effects on the nervous system have not been identified [44].

The administration of the 3DMAOM to the mouse did not alter the activities of ALP in the brain, suggesting that the plasma membrane integrity of the neurons was not adversely affected. Pregnane glycoside-rich extract normalized the ALP levels in streptozotocin-induced diabetic rats [45]; hence, rather than disruptive potentials, the 3DMAOM may have protective effects on the mouse’s brain. ALP is a widely distributed cell surface protein, which is also found in the brain and is soluble in various body fluids, including blood [46], and its elevation may be physiological or pathological. The physiological role of these enzymes is not entirely clear, but production increases in tissues undergoing metabolic stimulation [47]. Brain and plasma ALP activity increase has been linked to neurodegenerative diseases [46]. Moreover, ALP activity is a marker for plasma membrane and endoplasmic reticulum integrity [48].

In this study, CK activity was not altered in the brains of mice administered the 3DMAOM (**Figure 4**). This suggests that the 3DMAOM may not adversely affect energy homeostasis in the brain. The reported cellular energy modulatory effect of pregnane glycoside may not be via the brain CK-mediated pathway [49]. Brain-type CK deficiency is reportedly coupled to loss of function in neural cell circuits, altered bone-remodeling by osteoclasts, and complement-mediated phagocytotic activity of macrophages, processes sharing a dependence on actomyosin dynamics [50]. The integrity of the neural cell circuits appears to be maintained in the mice following the administration of 3DMAOM.

When the body is exposed to toxic compounds, AChE associates with these compounds, accumulating acetylcholine at the postsynaptic cleft sites [51]. The decreased AChE activity increases acetylcholine concentration in the synaptic cleft, resulting in neurological failures in impulse transmission [51]. Determination of the AChE activities in blood and tissue is useful in determining the exposure status of AChE inhibitors and detecting the toxicity of these compounds [52]. In this study, therefore, AChE activity in the brain was not significantly altered (**Figure 4**), suggesting that the 3DMAOM may not adversely affect the deactivation of acetylcholine after binding to its receptor in postsynaptic neurons during impulse transmission in cholinergic neurons and at neuromuscular junctions. This contradicts previous studies that classified pregnane glycosides from plant origin as exhibiting anti-AChE activities [52–53]. However, these studies are mechanistic in-vitro explorations, which may not translate to in-vivo effects. The specific carbon-carbon configuration in these various classes of pregnane glycoside may also contribute to various effects on biological parameters.

The activities of Na^+^/K^+^-, Ca^2+^/Mg^2+^-, and Mg2+-ATPases were not significantly affected (Figure 3), suggesting that administering 3DMAOM to mice may not adversely affect the electron gradient across the neuronal or nephron membrane. Thus, electrical signaling was maintained across the neurons or nephron membrane bilayer. 3DMAOM may not adversely affect the release of neurotransmitters at nerve endings and the subsequent transmission of impulses across the chemical synapse. Also, the crucial role of Na^+^/K^+^-ATPase in the brush border membrane in pumping Na^+^ back to the extracellular fluid was not compromised. Pregnane glycosides showed weak binding to Na^+^/K^+^-ATPase receptor which may explain the inability to significantly affect their activity in this study [55], could also be speculated to have similar effects to the other ATPases investigated. Toxic compounds targeting the biological cell first overcome the fence the biological membranes mount. Membranes are cellular boundaries controlling the molecular traffic across the boundary, dictating what goes in and out. They are central to the communication between cells and signal transduction pathways. Membrane receptors serve as the main targets to recognize specific ligands selectively, which can trigger a cascade of functional cell responses [56]. Na^+^/ K^+^-ATPase, Ca^2+^/Mg^2+^-ATPase, and Mg^2+^-ATPase are membrane-bound ion pumps that establish and maintain ionic gradients across the plasma membrane in all animal cells, using the free energy resulting from the hydrolysis of intracellular adenosine triphosphate (ATP) [57]. Due to the great importance of Na^+^/K^+^-ATPase in the maintenance of neuronal resting membrane potentials and propagation of neuronal impulses, the malfunction of this enzyme has been associated with neuronal hyperexcitability, cellular depolarization, and swelling [58]. Alteration in the activities of these ATPases by actions of toxic chemicals has been reported to cause cell injury and death of neurons, the basic events of cerebral ischemia in epilepsy, and various neurodegenerative disorders [41–42].

The plasma concentrations of Na^+^, K^+^, Cl^-^, and PO_4_^3-^ in mice administered the 3DMAOM were not altered in all the groups compared to controls . This suggests that the osmoregulatory function of the kidney with respect to these electrolytes remained uncompromised. However, the plasma concentration of Ca^2+^ was reduced at all 3DMAOM doses. This observation suggests that 3DMAOM may threaten Ca^2+^ homeostasis by possibly reducing its absorption in the intestine, a process normally mediated by 1, 25-dihydroxy vitamin D_3_, or by enhancing its excretion via reducing its reabsorption in the kidney. These may lead to muscle spasms, cramps, neuromuscular irritability, cognitive impairment, and electrocardiographic changes that mimic myocardial infarction or heart failure [61]. As the kidneys play a pivotal role in the regulation of body fluids, electrolytes, and acid-base balance, loss of kidney function predictably results in multiple derangement of a variety of electrolyte and acid-base, including hyperkalemia, metabolic acidosis, and hyperphosphatemia which, in turn, lead to serious complications including muscle wasting, bone-mineral disorder, vascular calcification, and mortality [62]. *G. latifolium* leaves extract significantly increased the antioxidant capacity in the kidney of streptozotocin-induced diabetic rat [63] indicative of their protective potential. However, 3DMAOM in the *G. latifolium* leaves may not be contributing to the maintaince of Ca^2+^ balance.

The activities of both alkaline phosphatase and GGT were not significantly changed in the kidneys of mice administered the 3DMAOM in this study. This suggests that the compound may not adversely affect the structural integrity of the plasma membrane and endoplasmic reticulum of the kidney cells and the role of GGT in intracellular glutathione metabolism, which is important in protecting intracellular protein thiol groups. In diabetic rats, the *G. latifolium* leaf extract normalized the levels of GGT [64], suggesting its protective effects instead of adverse changes. The relevant application of determination of enzyme activities in tissues and body fluids in clinical investigation and diagnosis cannot be overemphasized. Without structural or morphological alteration, tissue enzyme assay can indicate tissue cellular damage, giving insight into the site assaulted by the administered chemical or the organism exposed to [65]. Membrane-bound alkaline phosphatase and GGT serve as indices of renal injury [66].

The kidneys also play a vital role in excreting waste products such as urea, creatinine, and uric acid. Creatinine is the by-product of creatine phosphate in muscle, and it is produced at a constant rate by the body. For the most part, the kidney clears creatinine from the blood entirely. Urea is a nitrogen-containing compound formed in the liver as the end product of protein metabolism and the urea cycle. A large portion of urea is eliminated via the kidneys. Uric acid is the final oxidation product of purine metabolism and is excreted via the kidneys. Uric acid has been implicated in the pathophysiology of renal disease. Thus, plasma creatinine, urea, and uric acid levels assess renal sufficiency, especially glomerular filtration. Significant reduction in the glomerular filtration rate or obstruction of urine elimination results in increased blood creatinine, urea, and uric acid levels [47–48]. In this study, the plasma levels of urea and creatinine were unaltered compared to the control, suggesting that the glomerular filtration in the kidney was not compromised. In contrast, however, the plasma concentration of uric acid was significantly decreased at 2 mg/kg body weight of the 3DMAOM compared to controls, suggesting a reduction in the production of uric acid rather than insufficiency of glomerular filtration.

Finally, multivariate analysis was conducted, as shown in **Figure 5**. PCA is an unsupervised statistical method that involves reducing the size of principle components to examine changes among sample groups, and it is frequently utilized for exploratory data analysis [69]. **Figure 5A** displays the biplot of three principal components (PC1, PC2, and PC3), which account for >90% of the overall variability, indicating that the methods were successful [70,71,72]. PC1 accounts for the highest sample variation at 46.38%, while PC3 accounts for the lowest at 15.95%. PC1 primarily explains the most closely associated parameters, while PC2 and PC3 describe less interrelated parameters. In the biplot (**Figure 5A**), the 3DMA at 1, 2, and 4 mg/kg were located on the negative side of PC1, whereas 5% DMSO was situated on the positive side of PC2 showing the toxicity of 3DMA can be categorized based on their concentration level. Figure 5A shows that 3DMA at 1 shares similar characteristics with the other treatments as its confidence ellipse encompasses another dosage on the same PC 1. **Table S1** in the supporting information shows that PC1 is influenced by factors such as PCa, PUA, KGG, KAL, and KAL, with specific Eigenvector coefficients assigned to each. PC2 had the highest eigenvector coefficients for KMg (0.31771), BNa (-0.3795), BCK (0.3032), and KCa (0.3677). **Figure 5B** displays the results of hierarchical cluster analysis (HCA), revealing five separate clusters in a dendrogram, allowing for accurate classification. Cluster A strongly correlates BT, PK, and Plasma Ca^2+^ (PCa). Cluster B strongly correlates BNa, KTP, PCT, PUA, and KGG, whereas cluster E strongly correlates BA, KNa, Pur, KAL, and PPO. The results were consistent with the PCA findings.

**Figure 5.**
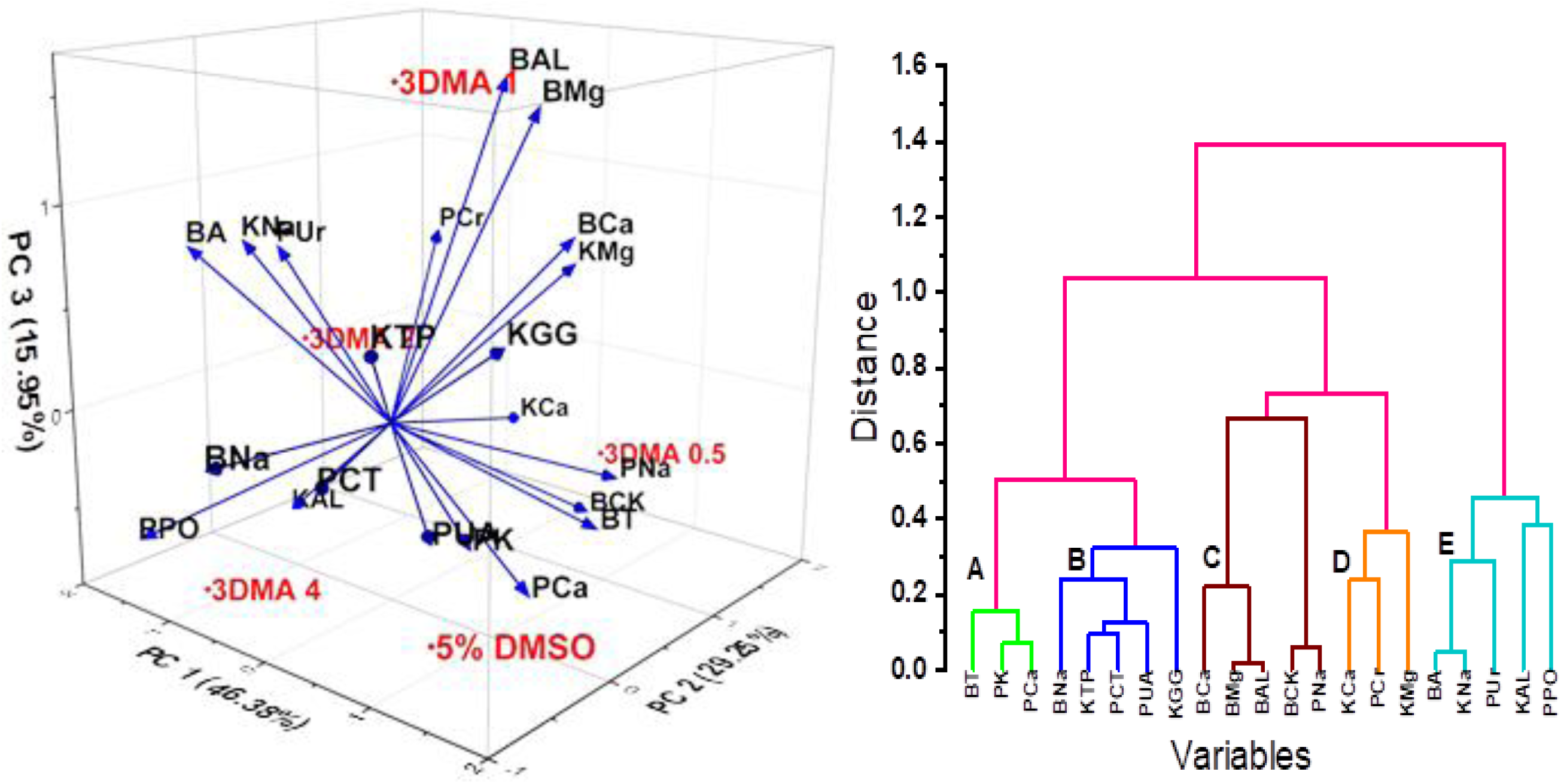
Multivariate illustrations of the effects of 3-*O*-[6-deoxy-3-*O*-methyl-β-D-allopyranosyl-(1→4)-β-D-oleandropyranosyl] -17β–marsdenin on brain and kidney indices in mice. **(A)** 3-dimensional biplot from principal component analysis (**B**) Hierarchical cluster analysis for various concentrations of 3-*O*-[6-deoxy-3-*O*-methyl-β-D-allopyranosyl-(1→4)-β-D-oleandropyranosyl. **Note.** BT, Brain Total Protein; BA, Brain AChE; BCa, Brain CaMgATPase; BMg, Brain MgATPase; BNa, Brain NaKATPase; BAL, Brain ALP; BCK, Brain CK; KTP, Kidney Total Protein; KCa, Kidney CaMgATPase; KMg, Kidney MgATPase; KNa, Kidney NaKATPase; KAL, Kidney ALP; KGG, Kidney GGT; PCr, Plasma Creatinine; PUr, Plasma Urea; PUA, Plasma Uric Acid; PNa, Plasma Na^+^; PCT, Plasma Cl^-^; PK, Plasma K^+^; PCa, Plasma Ca^2+^; PPO, Plasma PO_4_-.

## 5. Conclusion

Overall, the findings of this study suggest that 3DMAOM isolated from *Gongronema latifolium* leaf may be generally safe to the brain and kidney; however, it may adversely affect calcium ion homeostasis and possibly their role in nerve impulse transmission at chemical synapses. Therefore, further studies are recommended to delineate the mechanisms by which the compound causes derangement of plasma Ca^2+^ concentration, investigate the effects of other routes of administration of the compound, and chemically modify the structure of the compound to amend the observed electron impairment.

## List of Abbreviation

3DMAOM: 3-*O*-[6-deoxy-3-*O*-methyl-β-D-allopyranosyl-(1→4)-β-D-oleandropyranosyl]-17β–marsdenin
DMSO: Dimethyl sulphoxide
ALP: Alkaline phosphatase
CK: Creatine kinase
GGT: gamma-glutamyltransferase
AChE: Acetylcholinesterase
SDS: Sodium Dodecyl Sulfate

---

**Table S1.**
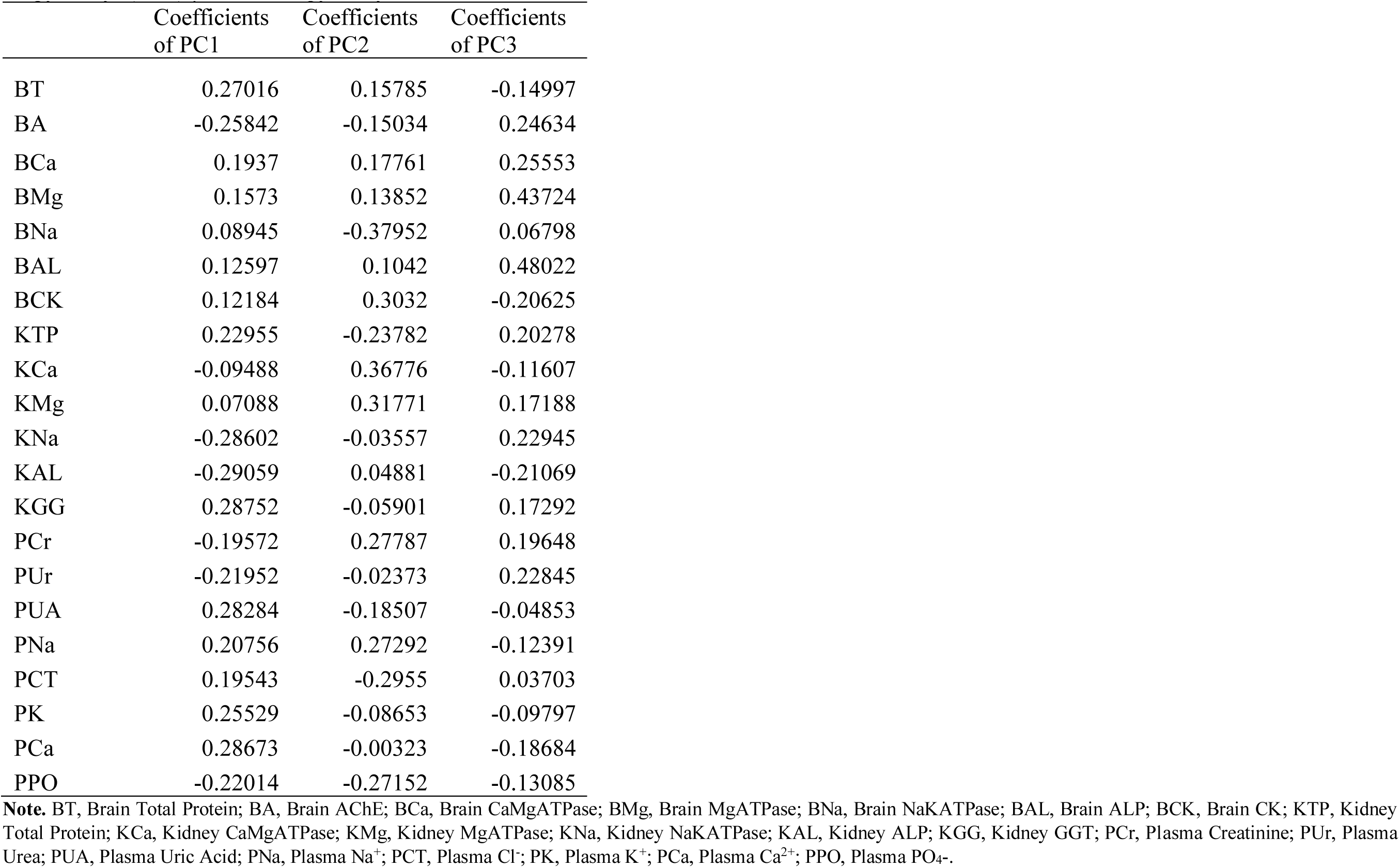
Coefficients of the eigenvectors were taken from different principal components analysis for various concentrations of 3-*O*-[6-deoxy-3-*O*-methyl-β-D-allopyranosyl-(1→4)-β-D-oleandropyranosyl.

