## Supplementary material for "Toxicity assessment of 3-*O*-[6-deoxy-3-*O*-methyl-β-D-allopyranosyl-(1→4)-β-D-oleandropyranosyl]-17β–marsdenin isolated from *Gongronema latifolium* leaf on selected brain and kidney function indices in mice": Table S1

### Supplementary data.

**Table S1**. Coefficients of the eigenvectors were taken from different principal components analysis for various concentrations of 3-*O*-[6-deoxy-3-*O*-methyl-β-D-allopyranosyl-(1→4)-β-D-oleandropyranosyl.

| | Coefficients of PC1 | Coefficients of PC2 | Coefficients of PC3 |
|---|---|---|---|
| BT | 0.27016 | 0.15785 | -0.14997 |
| BA | -0.25842 | -0.15034 | 0.24634 |
| BCa | 0.1937 | 0.17761 | 0.25553 |
| BMg | 0.1573 | 0.13852 | 0.43724 |
| BNa | 0.08945 | -0.37952 | 0.06798 |
| BAL | 0.12597 | 0.1042 | 0.48022 |
| BCK | 0.12184 | 0.3032 | -0.20625 |
| KTP | 0.22955 | -0.23782 | 0.20278 |
| KCa | -0.09488 | 0.36776 | -0.11607 |
| KMg | 0.07088 | 0.31771 | 0.17188 |
| KNa | -0.28602 | -0.03557 | 0.22945 |
| KAL | -0.29059 | 0.04881 | -0.21069 |
| KGG | 0.28752 | -0.05901 | 0.17292 |
| PCr | -0.19572 | 0.27787 | 0.19648 |
| PUr | -0.21952 | -0.02373 | 0.22845 |
| PUA | 0.28284 | -0.18507 | -0.04853 |
| PNa | 0.20756 | 0.27292 | -0.12391 |
| PCT | 0.19543 | -0.2955 | 0.03703 |
| PK | 0.25529 | -0.08653 | -0.09797 |
| PCa | 0.28673 | -0.00323 | -0.18684 |
| PPO | -0.22014 | -0.27152 | -0.13085 |

**Note.** BT, Brain Total Protein; BA, Brain AChE; BCa, Brain CaMgATPase; BMg, Brain MgATPase; BNa, Brain NaKATPase; BAL, Brain ALP; BCK, Brain CK; KTP, Kidney Total Protein; KCa, Kidney CaMgATPase; KMg, Kidney MgATPase; KNa, Kidney NaKATPase; KAL, Kidney ALP; KGG, Kidney GGT; PCr, Plasma Creatinine; PUr, Plasma Urea; PUA, Plasma Uric Acid; PNa, Plasma $Na^+$; PCT, Plasma $Cl^-$; PK, Plasma $K^+$; PCa, Plasma $Ca^{2+}$; PPO, Plasma $PO_4$-.
